# GRAPE: Genomic Relatedness Detection Pipeline

**DOI:** 10.1101/2022.03.11.483988

**Authors:** Aleksandr Medvedev, Mikhail Lebedev, Andrew Ponomarev, Mikhail Kosaretskiy, Dmitriy Osipenko, Alexander Tischenko, Egor Kosaretskiy, Hui Wang, Dmitry Kolobkov, Vitalina Chamberlain-Evans, Ruslan Vakhitov, Pavel Nikonorov

## Abstract

Classifying the degree of relatedness between pairs of individuals has both scientific and commercial applications. As an example, GWAS may suffer from high rates of false positive results due to unrecognized population structure. This problem becomes especially relevant with recent increases in large-cohort studies. Accurate relationship classification is also required for genetic linkage analysis to identify disease-associated loci. Additionally, DNA relatives matching service is one of the leading drivers for the direct-to-consumer genetic testing market. Despite the availability of scientific and research information on the methods for determining kinship and the accessibility of relevant tools, the assembly of the pipeline, that stably operates on a real-world genotypic data, requires significant research and development resources. Currently, there is no open-source end-to-end solution for relatedness detection in genomic data, that is fast, reliable and accurate for both close and distant degrees of kinship, combines all the necessary processing steps to work on real data, and is ready for production integration. To address this, we developed GRAPE: Genomic RelAtedness detection PipelinE. It combines data preprocessing, identity-by-descent (IBD) segments detection, and accurate relationship estimation. The project uses software development best practices, as well as GA4GH standards and tools. Pipeline efficiency is demonstrated on both simulated and real-world datasets.

## Introduction

Distant relationship estimation has both scientific and commercial applications. *Scientific* applications may include the identification of monogenic (single gene) Mendelian diseases [1], [2]. Also, relatedness detection can be used during data quality control for GWAS, since close relatives should be excluded to ensure that no pair of individuals is more closely related than second degree relatives [3], [4]. Otherwise, GWAS may suffer from high rates of false positive results. Potential *commercial* application of relationship estimation is utilized by direct-to-consumer genetic testing companies to find possible distant relatives for their customers [5]. The applicability of existent tools and pipelines to both applications is limited. Currently, there is no open-source end-to-end solution for relatedness detection in genomic data, that: (a) is reliable and accurate for both close and distant degrees; (b) includes all necessary processing steps to work with real data; (c) is flexible enough to be adapted for various applications; (d) is user-friendly and ready for production integration. Specifically, for commercial usage, the pipeline should deal with data heterogeneity and efficiently process newly added data samples. Samples can be genotyped with different chips and may have been passed thought different quality controls. Typical databases of genetic testing companies contain a lot of data from the previously used chips, which need to be combined together during the processing. Scientific database may also contains heterogeneous data. For example, UK Biobank dataset [6] contains samples which were genotyped using two different chips.

Driven by the idea of open and user-led innovation, we have developed GRAPE (Genomic RelAtedness detection PipelinE), the first open-source end-to-end solution for relatedness detection that is able successfully address the above mentioned difficulties. As a preliminary step, we comprehensively studied available approaches and software instruments for relatedness estimation from genotypes data, such as identity-by-descent (IBD) segments detection tools: GERMLINE [7], IBIS [8], RaPID [9], PhasedIBD [10]; relationship inference tools: DRUID [11], ERSA [12, 13], KING [14]; other tools, which may be required during data preprocessing steps, like algorithms of phasing (Eagle [15]) and genotype imputation (Minimac [16]). Then, we selected several perspective tools and joined them together into the user-friendly GRAPE pipeline.

GRAPE adapts the best practices for software development, including the Snakemake [17] workflow management system, Conda [18] virtual environments, Docker [19] containerisation, Funnel task execution service, which implements GA4GH Task Execution Schema [20], and CI/CD with automatic testing. The pipeline requires a single multi-sample VCF file as input and has a separate workflow for downloading of reference datasets and checking their consistency. IBD segments detection workflows of GRAPE can work with both phased and unphased data. As real-world datasets are often heterogeneous and inconsistent, GRAPE incorporates various data preprocessing and quality control (QC) options. GRAPE has a modular architecture that allows switching between tools and adjust tools parameters for better control of precision and recall levels. The pipeline also contains a simulation workflow with an in-depth evaluation of pipeline accuracy using simulated and reference data.

GRAPE can work like a standalone version as well as from the dedicated Docker container. Provided docker image contains all the dependencies installed, and highly recommended to use. We published GRAPE image in both Docker Hub and Dockstore [21] repositories to satisfy GA4GH [22] standards for sharing Docker-based tools. As a results we got a robust, reliable, and easy-to-use tool. Analysis for 10k samples with 600k SNPs requires half an hour, and about 22 hours have been required to process 100k samples dataset. Precision / recall analysis for GRAPE was performed using simulated datasets. We compared GRAPE with TRIBES [23], another open-source pipeline for relatedness detection, and showed the advantages of our solution in a sense of the precision / recall metrics.

## Methods

This section describes the input data, reference datasets, and the main pipeline steps. Scheme of the pipeline is presented in Figure 1. GRAPE takes single VCF file of individuals genotypes as input. Configuration of the pipeline is managed by `config.yaml` file, or via the parameters of the GRAPE launcher. Reference data should be previously downloaded and stored on a hard drive. After that, the pipeline performs relationship inference accordingly to one of the three possible workflows (described in a corresponding section below). The main pipeline steps are:

**Figure 1.**
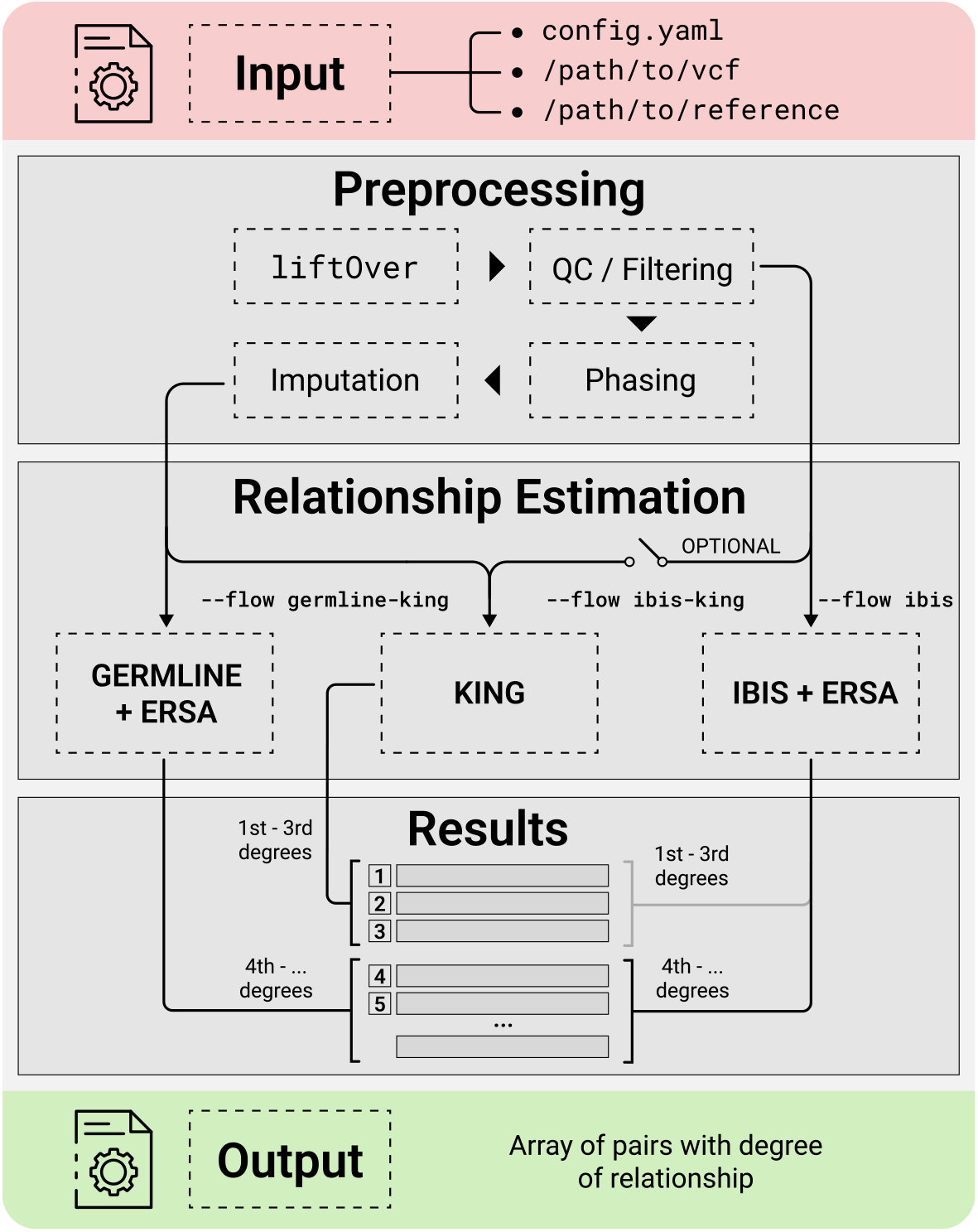
Scheme of the GRAPE pipeline.

1. Downloading of the reference datasets.
2. Quality control and data preprocessing.
3. Application of the relationship inference workflow.

GRAPE also includes simulation workflow to evaluate precision / recall metrics on the simulated data. To simulate artificial pedigrees, Ped-sim tool [24] is used, while unrelated founders for the simulation are taken from 1000 Genomes Project data.

There are two main approaches for relationship estimation implemented in GRAPE. The first one is based on allele frequencies calculation (KING [14]). This approach is optionally used for the first three degrees of relatedness, and for calculation of the kinship coefficients. The second approach relies on searching of pairwise identical-by-descent (IBD) segments. There are two options for this purpose based on two different tools: (a) IBIS [8] and (b) GERMLINE [7].

IBIS is a fast tool that can operate with unphased data. IBIS performs an IBD detection using homozygous SNPs only. It breaks the genome into windows of fixed length and searches for homozygous SNP mismatches for each genotype window. If there the number of mismatches does not exceed some predefined threshold, the window becomes a part of an IBD segment. In contrast, GERMLINE is slower and can work only with phased data, but under some circumstances it may produce higher precision results. Upon the pairwise IBD segments search is completed, relationship degrees are estimated using the ERSA tool [12].

### Downloading of the Reference Datasets

GRAPE requires various reference data to perform preprocessing, quality control, phasing, and imputation. Reference data is also required for the simulation workflow. In order to facilitate collection of reference data, we created a separate workflow that automates this step. It can be run by specifying `reference` command to the GRAPE pipeline launcher. The workflow downloads data, unpack it, and perform required post-processing procedures. If phasing and genotype imputation are required, one should also add additional flags `--phase` and `--impute` to the command. It affects the amount of downloaded data. If these flags are specified, the workflow downloads additional reference dataset to make phasing and imputation possible.

There is another options to download all required reference data as a single file. This file is prepared by us and preloaded on our side in the cloud. It can be done by specifying additional flag `--use-bundle` to the workflow. This way is faster, since all the post-processing procedures have been already performed. Reference files consist of three main groups.

- **Files for the preprocessing**. These files genetic recombination maps for mapping SNPs coordinates from base pairs (bp) to centimorgans (cM); files with the SNPs information from the 1000 Genomes Project for the SNPs quality control; reference genome of hg37 build; liftOver chain file.
- **Files for phasing and imputation**. Phasing is required as a preliminary step for GERMLINE tool, if input data is unphased. Upon the phasing is done, genotype imputation can be additionally applied. These files are space demanding and require a considerable amount of post-processing time.
- **Files for simulation**. These files include phased per-chromosome files from the 1000 Genomes Project; Affymetrix chip data that is used as a source of founders for the simulation; sex-specific recombination maps for better Ped-sim simulation results [24].

### Quality Control and Data Preprocessing

GRAPE have a versatile and configurable preprocessing workflow. One part of the preprocessing is required and must be performed before the relationship inference workflow. It is launched by the `preprocess` command of the GRAPE. Along with some necessary technical procedures, preprocessing includes the following steps.

- **[Required] SNPs quality control by minor allele frequency (MAF) and the missingness rate**. We discovered that blocks of rare SNPs with low MAF value in genotype arrays may produce false positive IBD segments. To address this problem, we filter SNPs by minor allele frequency. We remove SNPs with MAF value less than 0.02. Additionally, we remove multiallelic SNPs, insertions / deletions, and SNPs with the high missingness rate, because such SNPs are inconsistent with IBD detection tools.
- **[Required] Per-sample quality control, using missingness and heterozygosity**. Extensive testing revealed that samples with an unusually low level of heterozygosity could produce many false relatives matches among individuals. GRAPE excludes such samples from the analysis and creates a report file with the description of the exclusion reason.
- **[Required] Control for strands and SNP IDs mismatches**. During this step GRAPE fixes inconsistencies in strands and reference alleles.
- **[Optional] LiftOver from hg38 to hg37**. Currently GRAPE uses hg37 build version of the human genome reference. The pipeline supports input in hg38 and hg37 builds. One should specify the genome build version by the dedicated flag `--assembly` of the pipeline launcher. If hg38 build is selected (`--assembly hg38`), then GRAPE applies liftOver tool to the input data in order to match the hg37 reference assembly. This parameter should be also specified during the simulation workflow.
- **[Optional] Phasing and imputation**. GRAPE supports phasing and genotype imputation. GERMLINE IBD detection tool requires phased data. So, if input data is unphased, one should include phasing (`--phase` flag) into the preprocessing before running the GERMLINE workflow. If input data is highly *heterogeneous* in a sense of available SNPs positions, we recommend to include imputation procedure as well (`--impute`).
- **[Optional] Removal of imputed SNPs**. We found, that if input data is *homogeneous* in a sense of SNPs positions, the presence of imputed SNPs does not affect the overall IBD detection accuracy of the IBIS tool, but it significantly slows down the overall performance. For this particular case, when input data initially contains a lot of imputed SNPs, we recommend to remove them by specifying `--remove-imputation` flag to the GRAPE launcher. GRAPE removes all SNPs which are marked with `IMPUTED` flag in the input VCF file.

### GRAPE Workflows

There are three relationship inference workflows implemented in GRAPE. These workflows are activated by the `find` command of the launcher. Workflow selection is made by the `--flow` parameter.

1. **IBIS + ERSA**, `--flow ibis`. During this workflow IBD segments detection is performed by IBIS [8], and estimation of relationship degree is carried out by means of ERSA algorithm [12]. This is the fastest workflow.
2. **IBIS + ERSA & KING**, `--flow ibis-king`. KING [14] is a well-known method for the inference of close relationships. It’s fast and can work with unphased data. During this workflow, GRAPE uses KING tool for the first 3 degrees of relationships, and IBIS + ERSA approach for higher order degrees (see Figure 1). Comparison of evaluation time between IBIS + ERSA and IBIS + ERSA & KING workflows is presented in Figure 3.
3. **GERMLINE + ERSA & KING**, `--flow germline-king`. The workflow uses GERMLINE for IBD segments detection. KING is used to identify relationship for the first 3 degrees, and ERSA algorithm is used for higher order degrees. This workflow was added to GRAPE mainly for the case, when input data is already phased and accurately preprocessed.

### Pedigree Simulation

We added a simulation workflow into GRAPE to perform a precision / recall analysis of the pipeline. It’s accessible by `simulate` command of the pipeline launcher and incorporates the following steps: (1) pedigree simulation with unrelated founders; here we use the Ped-sim simulation package [24]; (2) relatedness degrees estimation; (3) comparison between true and estimated degrees. The source dataset for the simulation is taken from CEU population data of 1000 Genomes Project. As CEU data consists of trios, we picked no more than one member of each trio as a founder. We also ran GRAPE on selected individuals to remove all cryptic relationships up to the 6th degree. Then, we randomly assigned sex to each individual and used sex-specific genetic maps to take into account the differences in recombination rates between men and women [25]. Results of our precision / recall analysis for the simulated datasets are presented in the corresponding Results section.

### IBD Segments Weighing

Distribution of IBD segments among non-related (background) individuals within a population may be quite heterogeneous [13]. There may exist genome regions with extremely high rates of overall matching, which are not inherited from the recent common ancestors [26]. Instead, these regions more likely reflect other demographic factors of the population. The implication is that IBD segments detected in such regions are expected to be less useful for estimating recent relationships. Moreover, such regions potentially prone to false-positive IBD segments.

GRAPE can use two different approaches to address this issue. The first one is based on genome regions exclusion mask, wherein some genome regions are completely excluded from the consideration. This approach was proposed by authors of the ERSA algorithm, see [13]. The mask was computed based on whole-genome sequencing data for European individuals. The computed mask is built-in into ERSA 1.0 algorithm and is used by GRAPE by default.

The second approach is based on the so-called IBD segments weighing. This idea reminds one proposed in the Ancestry DNA Matching White Paper [27] (see description of their Timber algorithm there). The key idea is to down-weight IBD segment, i.e. reduce the IBD segment length, if the segment cross regions with high rate of matching. The approach can be briefly described as follows. At first, one should evaluate IBD segments {*B*_*j*_} for some background population. After that, we break chromosomes into windows *W*_*i*_ of fixed length and compute total overlap length *c*_*i*_ between found IBD segments {*B*_*j*_} and each window *W*_*i*_, *c*_*i*_ = Σ_*j*_ |*W*_*ij*_|, where *W*_*i j*_ is an overlap between the segment *B*_*j*_ and the window *W*_*i*_. Obtained overlap lengths are transformed into weights *w*_*i*_ which are assigned to the windows, *w*_*i*_ = *f* (*c*_*i*_) ∈ [0; 1]. Here *f* is a weighing function that reflect the following heuristic: if the overlap length for a window *W*_*i*_ is a relative outliers in the distribution of overlap lengths, then value of *f* (*c*_*i*_) is close to 0, otherwise it’s close to 1. Given that weights for all windows are computed, for each IBD segment *G* one can compute its weighted length |*G*|_w_ using the formula

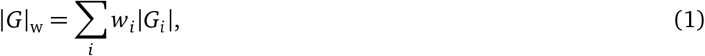

where |*G*_*i*_| denotes the overlap length between IBD segment *G* and a window *W*_*i*_. Weighted lengths of IBD segments then are used by the ERSA algorithm (while the ERSA mask is disabled).

GRAPE provides an ability to compute the weight mask from the VCF file with presumably unrelated individuals. It breaks each chromosome into 1cM windows to compute overlaps. After that, GRAPE detects outliers among overlap lengths by means of Minimum Covariance Determinant (MCD) algorithm [28], and then determines outliers upper bound *h*. This upper bound is used to compute weights, *w*_*i*_ = *h*/*c*_*i*_, if *c*_*i*_ > *h*; and *w*_*i*_ = 1 otherwise. Computed mask can be further used as a parameter for relationship inference workflow (see the `--weight-mask` parameter).

As an example, Figure 2 depicts the weight mask computed for the individuals of East Asian Ancestry taken from 1000 Genomes Project. To detect IBD segments, IBIS workflow was used with parameters `--ibis-seg-len 5`, `--ibis-min-snp 400`. High matching regions (*w*_*i*_ ≈ 0) are marked with the blue color. For comparison, ERSA 1.0 masked regions are depicted with the pick hatching.

**Figure 2.**
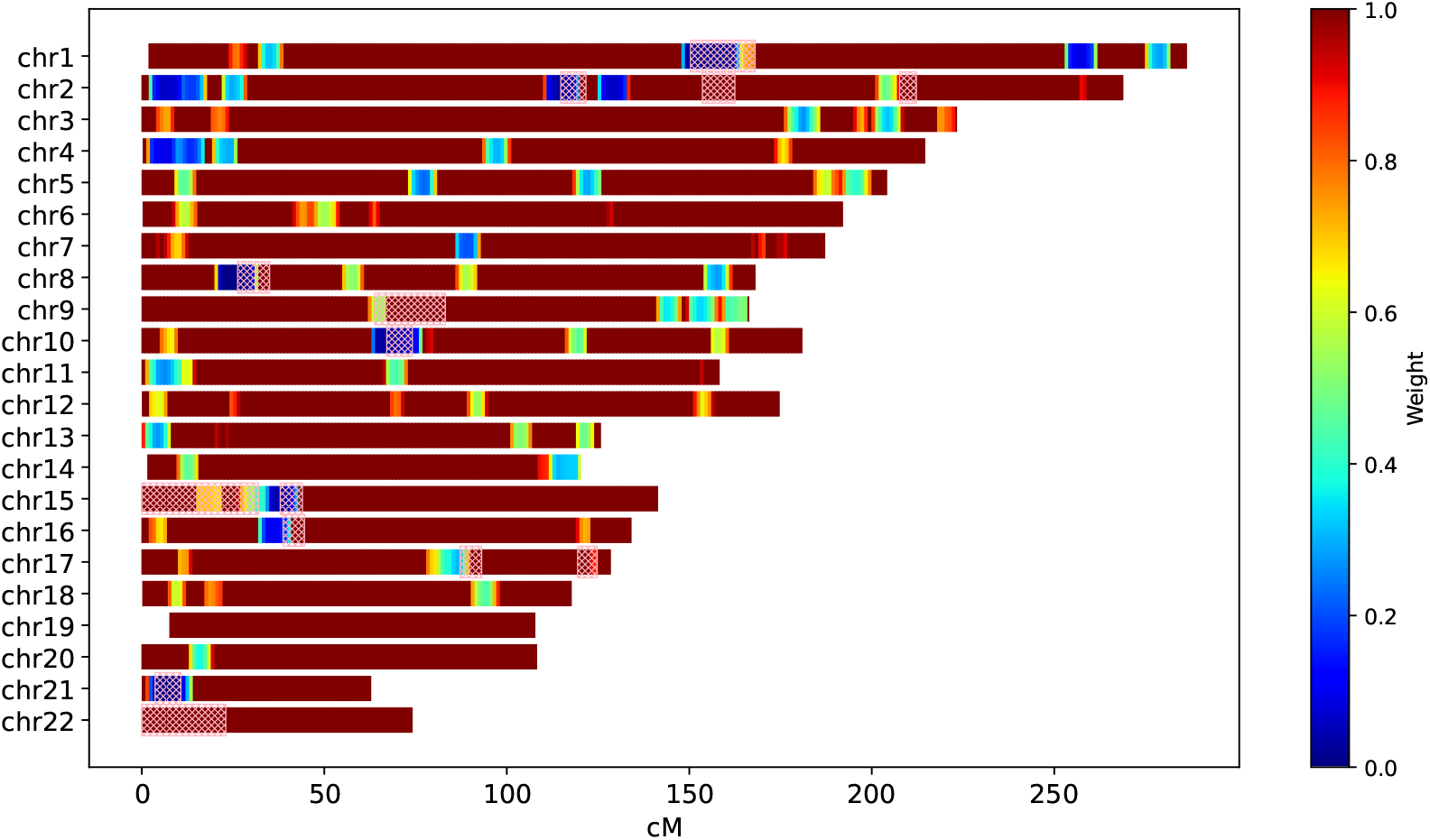
Weight mask computed for the individuals of East Asian Ancestry taken from 1000 Genomes Project; compared to ERSA 1.0 masked regions (pink hatching). IBIS workflow parameters: `--ibis-seg-len 5`, `--ibis-min-snp 400`.

### Operation

GRAPE can be run inside a Docker container. This way is recommended. Another option is to run the pipeline from scratch with all the dependencies being pre-installed. We successfully ran the pipeline on Ubuntu 18.04 with 8 CPUs and 32 GB of RAM to evaluate the performance on huge datasets with up to 100k samples and millions of SNPs.

### Resource Allocation

GRAPE can utilize multiple cores. For that, one should specify the cores number via the `--cores` parameter. Default number of cores equals to the total number of available CPUs minus 1.

### Execution by Scheduler

The pipeline can be run using Funnel [29], a lightweight task scheduler that implements Task Execution Schema [20] developed by GA4GH [22]. The scheduler can work in various environments, from a regular virtual machines to Kubernetes cluster with the support of resource quotas. We provide several examples of the task specifications for Funnel.

Each sample represents a JSON file with the task description. These files are available in the GRAPE GitHub repository within the corresponding `funnel` subfolder.

### Performance

The fastest IBIS + ERSA (`--flow ibis`) relatedness inference workflow takes about 22 hours to process 100k individuals dataset, see Figure 3. IBIS tool has a quadratic time complexity with respect to the total number of individuals. The addition of KING (`--flow ibis-king`) increases the total running time by roughly 50%. Performance analysis confirmed that IBIS is a simple and efficient tool. It allows the pipeline to process hundred of thousands of individuals in a reasonable amount of time.

**Figure 3.**
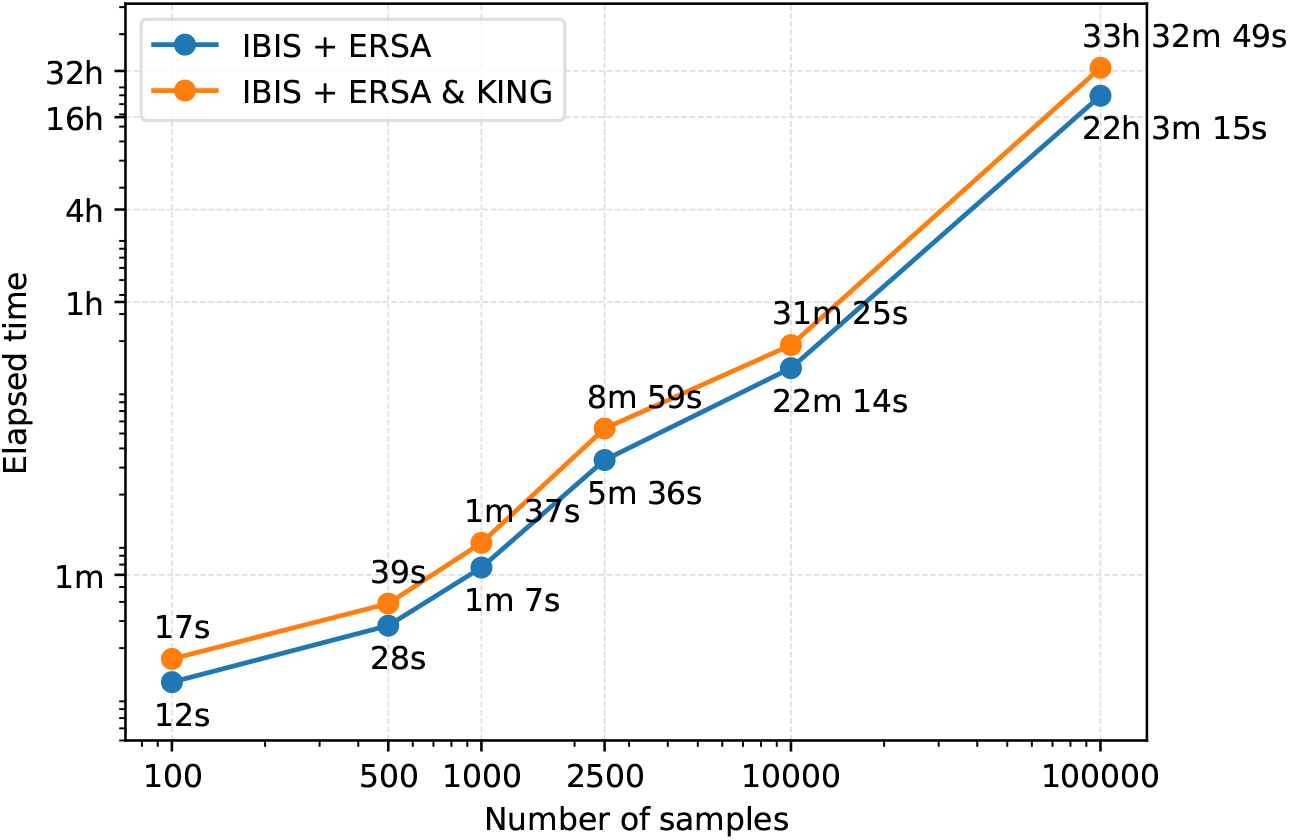
Performance comparison between IBIS + ERSA and IBIS + ERSA & KING workflows. GRAPE was evaluated on a machine with 8 CPU cores and 32 GB of RAM. Both axes are in logarithmic scale.

## Use Cases

This section gives examples of the GRAPE pipeline commands which have to be run to infer relationships, or to evaluate precision / recall metrics on a simulated dataset.

### Relationship Inference with IBIS + ERSA

As the first step, reference data must be downloaded. We suppose that reference data to be stored in `/media/ref` directory. Here and below we specify the `--real-run` flag. Without this flag GRAPE performs a dry-run.

**Listing 1**. Reference downloading for the IBIS + ERSA workflow

~~~
docker run --rm -it -v /media:/media
    genx_relatives:latest launcher.py reference --ref-directory /media/ref --real-run
~~~

As the second step, preprocessing is performed. We suppose that the input file is located at `/media/input.vcf.gz`, and it’s in hg38 build. Input file location specified by the flag `--vcf-file`. GRAPE working directory is `/media/data`. It’s specified by the `--directory` flag.

**Listing 2**. Preprocessing for the IBIS + ERSA workflow

~~~
docker run --rm -it -v /media:/media
    genx_relatives:latest launcher.py preprocess --ref-directory /media/ref
       --vcf-file /media/input.vcf.gz --directory /media/data --assembly hg38 --real-run
~~~

The third step is the relationship inference. It is launched with the the `find` command. We use IBIS + ERSA workflow that corresponds to the `ibis` value of the `--flow` parameter (default).

**Listing 3**. Relationship inference with the IBIS + ERSA workflow

~~~
docker run --rm -it -v /media:/media
    genx_relatives:latest launcher.py find --flow ibis --ref-directory /media/ref
       --directory /media/data --ibis-seg-len 7 --ibis-min-snp 500
       --zero-seg-count 0.5 --zero-seg-len 5.0 --alpha 0.01 --real-run
~~~

GRAPE has an ability to specify additional parameters to the ERSA and IBIS algorithms to control the sensitivity and the false positive rate. These parameters are described below. Default GRAPE parameters are quite conservative. They provide a low false positive rate and low sensitivity in high (9+) degrees.

- [IBIS] `--ibis-seg-len`. Minimum length of the IBD segment to be found by IBIS. Higher values reduce false positive rate and give less distant matches (default = 7 cM).
- [IBIS] `--ibis-min-snp`. Minimum number of SNPs per IBD segment to be detected (default = 500 SNPs).
- [ERSA] `--zero-seg-count`. Mean number of shared segments for two unrelated individuals in the population. Smaller values tend to give more distant matches and increase the false positive rate (default = 0.5).
- [ERSA] `--zero-seg-len`. Average length of IBD segment for two unrelated individuals in the population. Smaller values tend to give more distant matches and increase the false positive rate (default = 5 cM).
- [ERSA] `--alpha`. ERSA significance level [12] (default = 0.01).

### Relationship Inference with IBIS + ERSA & KING

The first and the second steps are the same as for the previous use case. Third step is launched with specifying of the `ibis-king` flow parameter. For this case GRAPE performs an estimation of the first three degrees of relationship with KING. Other degrees are estimated with ERSA (see Figure 1). Resulted report of individual relationships for this case contains additional `king_degree` and `kinship` columns. KING algorithm has no additional parameters.

**Listing 4**. Relationship inference with the IBIS + ERSA & KING workflow

~~~
docker run --rm -it -v /media:/media
    genx_relatives:latest launcher.py find --flow ibis-king --ref-directory /media/ref
       --directory /media/data --ibis-seg-len 7 --ibis-min-snp 500
       --zero-seg-count 0.5 --zero-seg-len 5.0 --alpha 0.01 --real-run
~~~

### Relationship Inference with GERMLINE + ERSA & KING

At first, reference data must be downloaded. GERMLINE tool works with phased data only. So, if input data is unphased, one should download additional reference dataset to perform phasing. For that purpose we use a reference panel from ∼1000 Genomes Project. This panel takes a considerable amount of disk space (∼ 25 GB), and requires significant time to download. To download this panel one should specify `--phase` flag while using the `reference` command.

**Listing 5**. Reference downloading

~~~
docker run --rm -it -v /media:/media
    genx_relatives:latest launcher.py reference
       --ref-directory /media/ref --phase --real-run
~~~

The second step is the data preprocessing. We use `--phase` flag to apply phasing procedure during this stage.

**Listing 6**. Preprocessing for the GERMLINE + ERSA & KING workflow

~~~
docker run --rm -it -v /media:/media
    genx_relatives:latest launcher.py preprocess --ref-directory /media/ref
       --vcf-file /media/input.vcf.gz --directory /media/data
       --assembly hg38 --phase --real-run
~~~

The third step is the relationship inference. One should specify `--flow germline` to use GERMLINE tool for the IBD segments detection. ERSA parameters are the same as for IBIS + ERSA workflow. Parameters of the GERMLINE are not configurable. Currently, we use the following sets of GERMLINE parameters: `-min_m 2.5`, `-err_hom 2`, `-err_het 1`. See GERMLINE documentation for the parameters description [7].

**Listing 7**. Relationship inference with GERMLINE + ERSA & KING workflow

~~~
docker run --rm -it -v /media:/media
    genx_relatives:latest launcher.py find --flow germline-king --ref-directory /media/ref
       --directory /media/data --zero-seg-count 0.5 --zero-seg-len 5.0
       --alpha 0.01 --real-run
~~~

### Evaluation of the IBIS + ERSA Workflow on a Simulated Dataset

To perform simulation, at the first step one should download *full* reference dataset by using the `reference` command with the `--phase` and `--impute` flags enabled. The second step is the simulation workflow. For that, one should use `simulate` command of the GRAPE launcher.

**Listing 8**. Evaluation of the IBIS + ERSA workflow on a simulated dataset

~~~
docker run --rm -it -v /media:/media
    genx_relatives:latest launcher.py simulate --flow ibis --ref-directory /media/ref
       --directory /media/data --sim-params-file params/relatives_average.def
       --sim-samples-file ceph_unrelated_all.tsv --assembly hg37 --real-run
~~~

Along with the parameters for preprocessing, ERSA and IBIS tools, simulation workflow has two additional parameters listed below.

- `--sim-params-file`. File with parameters of simulation for the Ped-sim tool. For more information see [24]. We have prepared several simulation parameters files, and stored them in the GRAPE repository on GitHub.
- `--sim-samples-file`. File with a list of individuals from 1000 Genomes Project which are used as founders for the simulations. One can choose `ceph_unrelated_all.tsv` (unrelated individuals from CEU population), or `all.tsv` (all individuals available in 1KGP).

The output files include a list of kinship matches found in simulated dataset, precision / recall plots, and a confusion matrix to compare the detected degrees of relationship with the true degrees. Detailed information on the computed metrics is presented in the Results section.

### Evaluation of GERMLINE + ERSA & KING Workflow on a Simulated Dataset

The first step is to download the reference dataset (see the previous section). The second step is to run `simulate` command with specifying `germline-king` flow and the `--phase` flag.

**Listing 9**. Evaluation of the GERMLINE + ERSA & KING workflow on a simulated dataset

~~~
docker run --rm -it -v /media:/media
    genx_relatives:latest launcher.py simulate --flow germline-king
       --ref-directory /media/ref --directory /media/data --assembly hg37 --phase
       --sim-params-file params/relatives_average.def
       --sim-samples-file ceph_unrelated_all.tsv --real-run
~~~

### Computation of the IBD Segments Weighing Mask

GRAPE has a dedicated command to compute the weight mask, `compute-weight-mask`. It performs IBD segments detection for the input VCF file and then analyse IBD segments distribution to compute the weight mask. The resulting files consist of a weight mask file in JSON format and a visualization of the mask (see Figure 2)

**Listing 10**. Computation of the IBD segments weighing mask

~~~
docker run --rm -it -v /media:/media
    genx_relatives:latest launcher.py compute-weight-mask
       --directory /media/background --assembly hg37
       --real-run --ibis-seg-len 5 --ibis-min-snp 400
~~~

For the example above the resulting files are store in the `/media/background/weight-mask/` directory.

### Usage of the IBD Segments Weighing Mask

To apply weight mask during the relatedness detection (`find` command), one should specify the mask file with the `--weight-mask` parameter. When used, the ERSA 1.0 exclusion mask is disabled.

**Listing 11**. Usage of the IBD segments weighing mask

~~~
docker run --rm -it -v /media:/media
   genx_relatives:latest launcher.py find --flow ibis --ref-directory /media/ref
      --weight-mask /media/background/weight-mask/mask.json
      --directory /media/data --assembly hg37
      --real-run --ibis-seg-len 5 --ibis-min-snp 400
~~~

## Results

To test the accuracy and flexibility of the pipeline, we performed an extensive testing on both real and simulated datasets. As a sanity check, we took Allen Ancient DNA Resource (AADR) dataset [30] and made sure that GRAPE does not produce any kinship matches between ancient and present-day individuals.

Next, we have run Khazar origins dataset [31] through the GRAPE. Khazar dataset contains 1770 samples from 106 Jewish and non-Jewish populations. The dataset contains significant amount of data from small homogeneous populations. The KING analysis was previously applied to this dataset by the authors [31] to identify close relatives up to the third degree. We applied GRAPE and found 1715 putative relationships. Most of them have a degree of 4+. This result highlights the fact, that total length of IBD segments in small homogeneous populations is several orders of magnitude higher then for the heterogeneous populations in Europe and Asia [32, 33]. This is an obvious obstacle for the relatedness detection. There is no method known to us to address this issue while using genotypic data obtained from SNP arrays. Whole genome sequencing has a potential to solve this problem, since rare mutations should break long IBD segments.

As for the data from genetic testing companies, GRAPE has been successfully applied to the database of 100k+ customers of Atlas Biomed [34], a company that provides direct-to-consumer genetic tests. During this test GRAPE was proven to be able to handle diverse data obtained from different chips, reference alignments, and other possible data inconsistencies.

### Precision / Recall Analysis on Simulated Datasets

Finally, GRAPE was evaluated on simulated datasets. We performed the simulation using unrelated founders from 1KGP Affymetrix genotype chip data. This chip contains approximately 900k SNPs. The simulation was carried out with the Ped-sim package. Using this tool, we produced several pedigree structures with 8 generations, and a maximum degree of relationship equals to 14. Then we joined all of the pedigrees into one dataset, and performed the precision / recall analysis of the GRAPE pipeline for the different flows. For each degree *i* of relationships we computed precision and recall metrics:

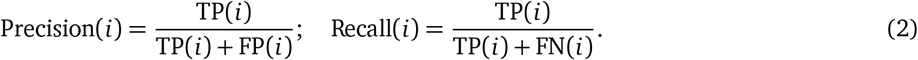

Here TP(*i*), FP(*i*), FN(*i*) are the numbers of true positive, false positive, and false negative relationship matches predicted for the degree *i*. In our analysis we used non-exact (fuzzy) interval metrics. For the 1st degree, we require an exact match. For the 2nd, 3rd, and 4th degrees, we allow a degree interval of ±1. For example, for the 2nd true degree we consider a predicted 3rd degree as a true positive match. For the 5th+ degrees, we use the ERSA confidence intervals which are typically 3-4 degrees wide. For 10th+ degrees, these intervals are 6-7 degrees wide. We also plot a confusion matrix for the predicted vs true degrees.

Results of the simulation for the IBIS + ERSA & KING workflow is presented in Figure 4. The following set of parameters was used: `--ibis-seg-len 7`, `--ibis-min-snp 500`, `--zero-seg-count 0.5`, `--zero-seg-len 5`, `--alpha 0.01`. Confusion matrix is presented on the right panel in Figure 4. There -1 stands for no-relationship. The pipeline shows recall above 90%+ for degrees from 1 to 5. It detects all relatives with 1-4 degrees. GRAPE found no false positive matches, i.e. it does not found any relationships among unrelated individuals, Recall(− 1) = 1. Precision is above 90% among all detected degrees. We set these parameters as default for this GRAPE workflow. These parameters are quite conservative, i.e. they provide high precision but low sensitivity.

**Figure 4.**
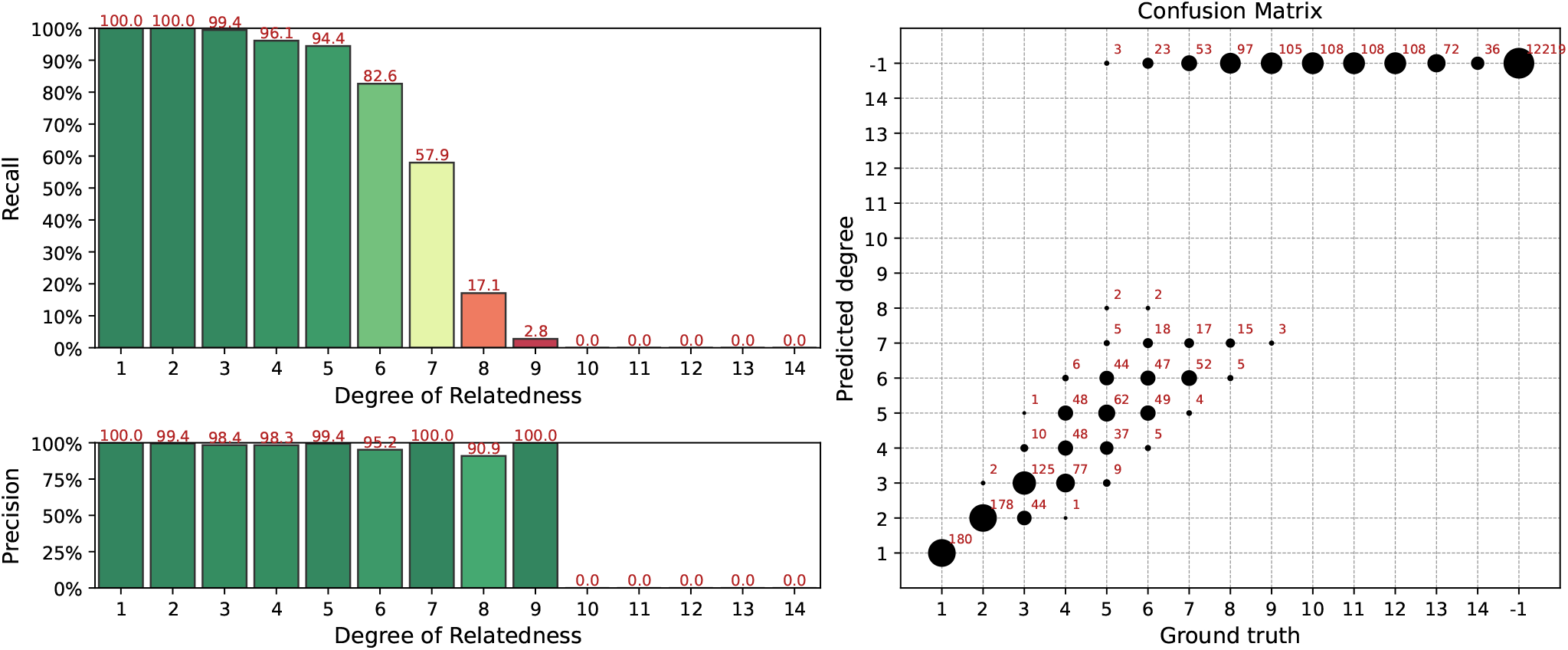
Interval (fuzzy) precision / recall (left panel) and confusion matrix (right panel) for the IBIS + ERSA & KING workflow, obtained for a simulated dataset. Parameters of the workflow: `--ibis-seg-len 7`, `--ibis-min-snp 500`, `--zero-seg-count 0.5`, `--zero-seg-len 5`, `--alpha 0.01`.

One can relax GRAPE parameters of get more relatedness matches for high degrees. On the other hand, the number of false positive matches increases as well. Figure 5 shows the simulation results for the IBIS + ERSA & KING workflow for slightly relaxed parameters: `--ibis-seg-len 5`, `--ibis-min-snp 400`, `--zero-seg-count 0.1`, `--zero-seg-len 5`, `--alpha 0.01`. Number of detected relationships increases significantly for higher (≥ 8) degrees. But false positive matches arise as well, i.e. Recall(−1) ≠ 1.

**Figure 5.**
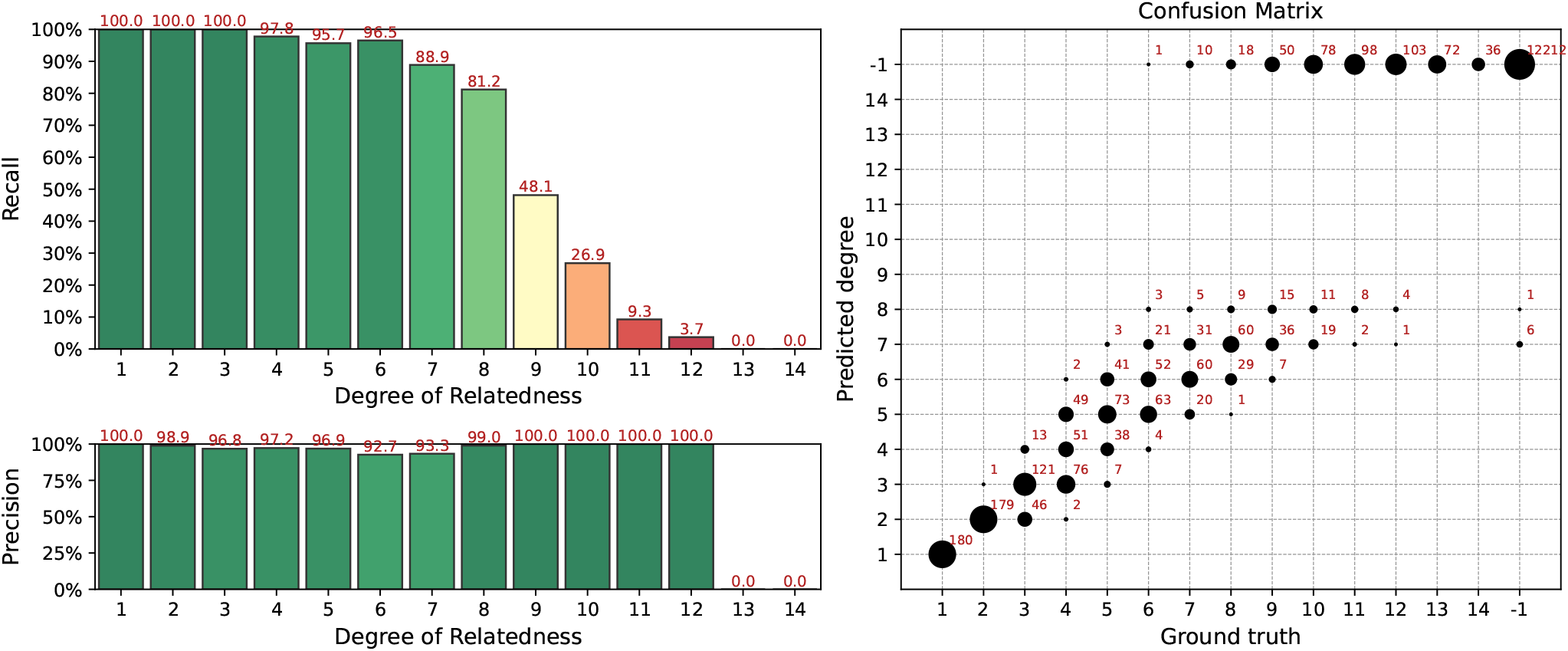
Interval (fuzzy) precision / recall (left panel) and confusion matrix (right panel) for the IBIS + ERSA & KING workflow, obtained for a simulated dataset. Parameters of the workflow: `--ibis-seg-len 5`, `--ibis-min-snp 400`, `--zero-seg-count 0.1`, `--zero-seg-len 5`, `--alpha 0.01`.

Simulation results for the GERMLINE + ERSA & KING workflow is presented in Figure 6. We used the same ERSA parameters as for Figure 4: `--zero-seg-count 0.5`, `--zero-seg-len 5`, `--alpha 0.01`. In comparison to Figure 4, one can see that GERMLINE slightly decrease recall for 6th and 7th degrees, but improves it for higher (≥ 8) degrees. Precision is above 95% among all detected degrees. No false positive matches were found, i.e. Recall(− 1) = 1. Our experiments showed that, in comparison to IBIS, GERMLINE is better suited for a careful analysis of relatively small cohorts with phased data. IBIS algorithm is less sensitive, but for segments with length above 7 cM, it produces the same results as GERMLINE, while working with unphased data.

**Figure 6.**
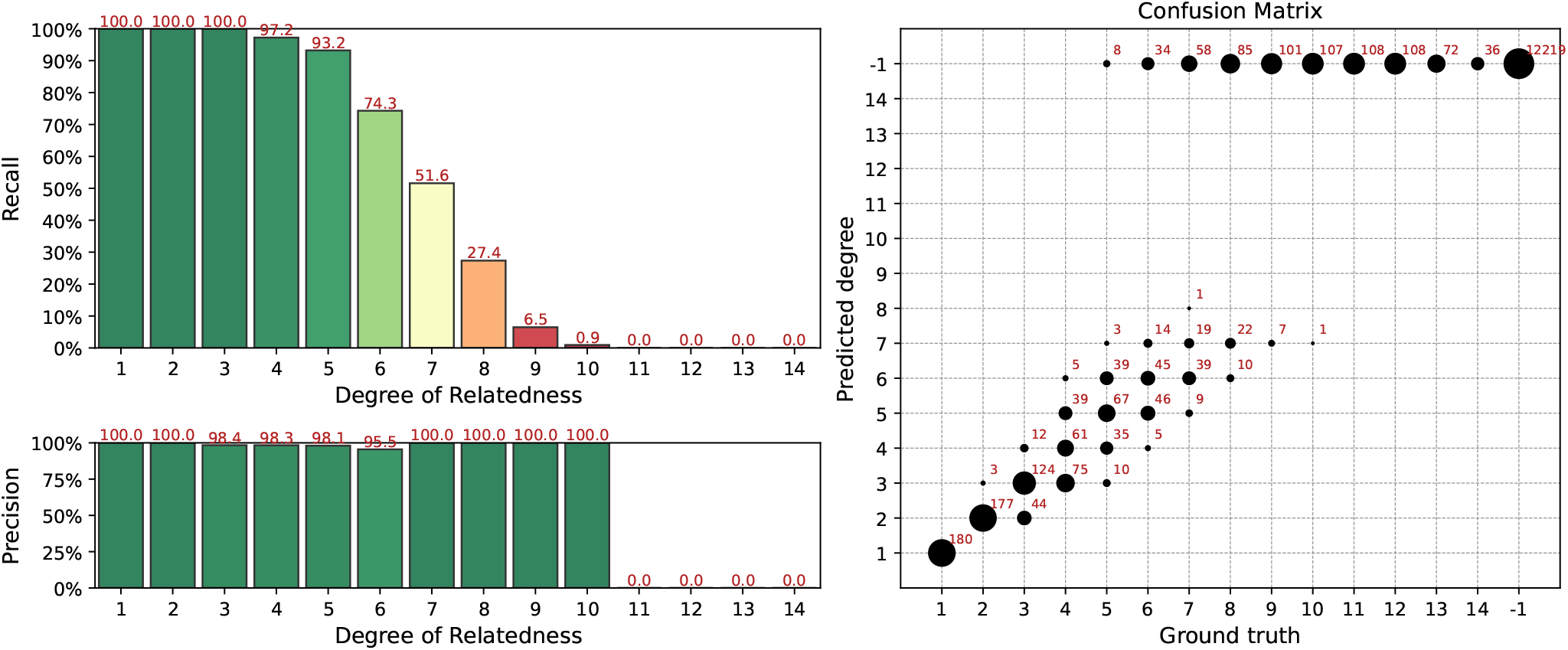
Interval (fuzzy) precision / recall (left panel) and confusion matrix (right panel) for the GERMLINE + ERSA & KING workflow, obtained for a simulated dataset. Parameters of the workflow: `--zero-seg-count 0.5`, `--zero-seg-len 5`, `--alpha 0.01`.

### Performance of the IBD Segments Weighing

Our simulation experiments showed that both weighing and exclusion approaches reduce false-positive rate and significantly improves the overall performance of the pipeline. By the way, weighing mask can be better adapted to specific ancestries, and after additional parameters tuning may slightly outperform ERSA 1.0 approach. In Figure 7 comparison between between the ERSA 1.0 exclusion mask and the GRAPE weight mask is presented for a simulated dataset with the founders of East Asian Ancestry from 1KGP. IBIS workflow was used. Weight mask was taken from Figure 2. Parameter `--zero-seg-count` was varied while using weight mask to achieve the same level of precision. One can see, that with approximately the same precision, weighing approach gives several percentages higher recall.

**Figure 7.**
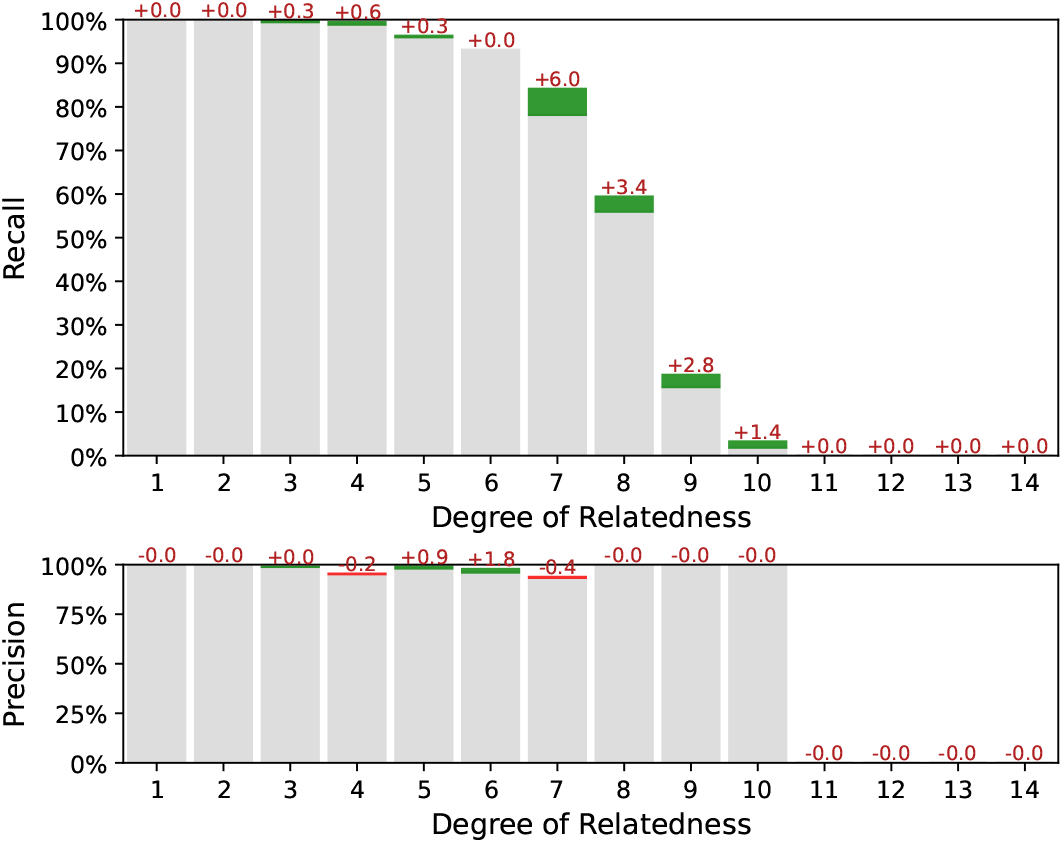
Interval (fuzzy) precision / recall comparison between the ERSA 1.0 exclusion mask and the GRAPE weight mask. Metrics differences are marked with colors: green, if the metric has increased while using weighing; and red, if the metric has decreased. Common parameters for the both workflows: `--ibis-seg-len 5`, `--ibis-min-snp 400 --zero-seg-len 5`, `--alpha 0.01`. Parameter `--zero-seg-count` equals `0.1` while using weight mask, and `0.3` while using original ERSA 1.0 exclusion mask.

### Comparison with TRIBES

In the end, we compared GRAPE with TRIBES [23]. TRIBES is an earlier open-source pipeline for relatedness detection. The pipeline combines the GERMLINE algorithm for IBD segments detection and the calculation of the genome proportion with zero alleles inferred IBD (IBD0) for each pair to detect the relatedness. If the data is not phased, TRIBES provides an ability to phase data with the EAGLE tool. This part of TRIBES is similar to one of the corresponding GRAPE workflows of IBD segments detection. In contrast to GRAPE, TRIBES estimates degrees of relationship according to expected IBD0 segments proportion ranges. GRAPE uses the ERSA algorithm, which, to our knowledge, is a more advanced approach.

We have run TRIBES pipeline on the same simulated datasets. Since the simulated datasets contain unphased data, we also applied built-in phasing procedure from TRIBES. The result of the analysis is presented in Figure 8. TRIBES has demonstrated high detection power for distant relationships of up to the 12th degree. Given that many 13+ degree relatives do not share any IBD segments, this is near a theoretical limit. However, TRIBES produces a *huge* number of false positive matches, see confusion matrix in the right panel of the Figure 8. Since TRIBES lacks options that allow users to control false positives rates by varying pipeline parameters, it becomes a crucial drawback. This obstacle does not allow the TRIBES pipeline to be adapted for applications, where desired precision / recall rate may vary depending on different business or research objectives.

**Figure 8.**
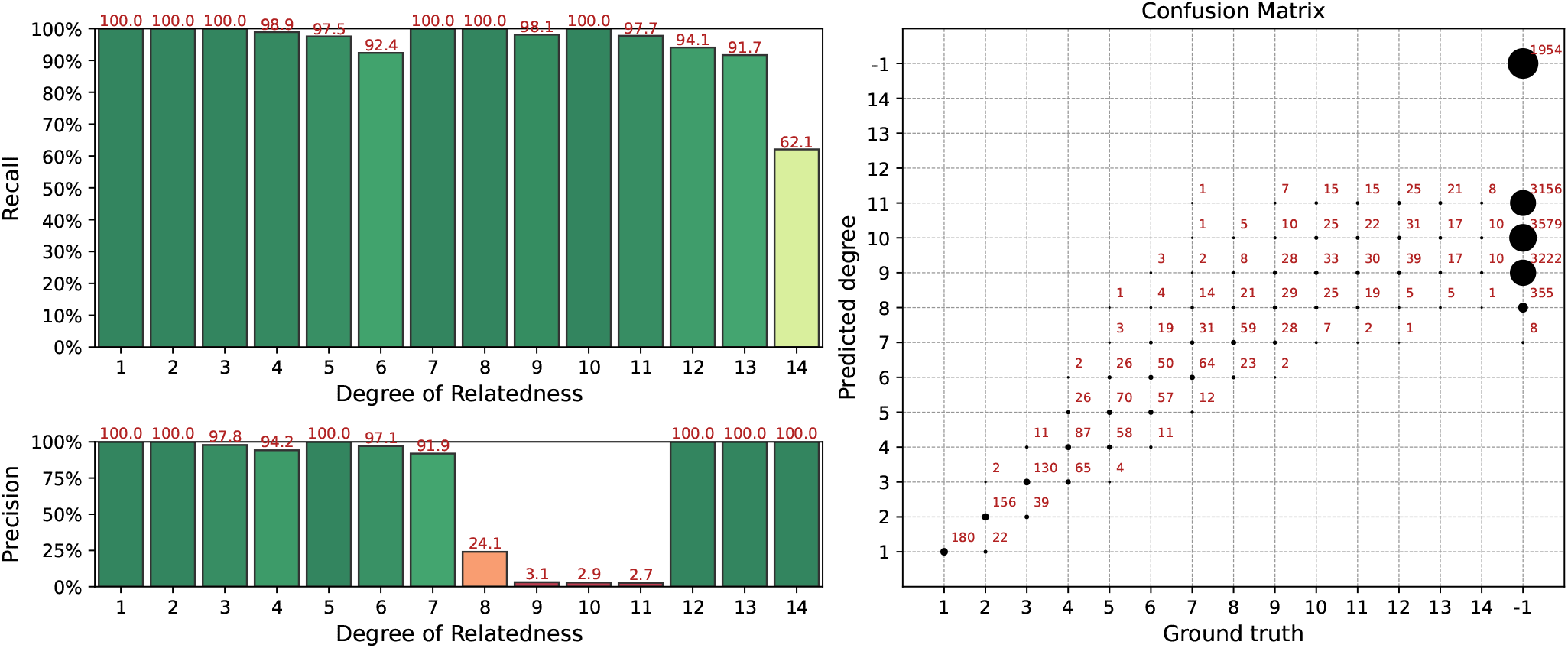
Interval (fuzzy) precision / recall (left panel) and confusion matrix (right panel) for the TRIBES pipeline results, obtained for a simulated dataset.

## Conclusions

In the current paper, we introduced GRAPE: genomic relatedness detection pipeline. We performed the careful selection of tools, and combined various preprocessing steps, IBD segments detection tools, and algorithms of estimations of relationship into a single pipeline. One of the possible workflows of the pipeline is based on IBIS tools and can work with unphased data. It’s suitable for the analysis of large cohorts in a relatively short time. Using this workflow GRAPE can perform relationship estimation among 100k samples in 22 hours. Another possible workflow is based on GERMLINE IBD segments detection tool and works with phased data. Our experiments showed that GERMLINE has the most detection power, while IBIS option is the fastest, easiest to use, and has sufficient accuracy for 1-8 degrees of relationship. Finally, we compared GRAPE with TRIBES, another relatedness detection pipeline. In contrast to GRAPE, TRIBES produce a huge number of false positive matches, requires phased data, and lacks of important preprocessing and evaluation options, that makes it impractical for various application. GRAPE is proved to be reliable and accurate tool for the analysis of close and distant degrees of kinship. It provides an ability to control a false positive rate, can work with heterogeneous data obtained from various chips, and is ready for production integration.

## Data availability

Public available datasets which are used in the article:

- 1000 Genomes Project [35]; URL: https://www.internationalgenome.org/data-portal/data-collection/phase-3.
- Khazar Origin for Ashkenazi Jews [31]; URL: https://evolbio.ut.ee/khazar.
- Allen Ancient DNA Resource (AADR) [30]; URL: https://reich.hms.harvard.edu/allen-ancient-dna-resource-aadr-downloadable-genotypes-present-day-and-ancient-dna-data.

Atlas Biomed customers dataset cannot be published due to the privacy reasons.

## Software availability

GRAPE pipeline can be accessed from the following pubic available resources:

- GitHub: https://github.com/genxnetwork/grape.
- Docker Hub: https://hub.docker.com/r/genxnetwork/grape.
- Dockstore: https://dockstore.org/organizations/GenX/collections/GRAPE. License: GPLv3.

## References

[1] Jennifer E. Posey, Anne H. O’Donnell-Luria, Jessica X. Chong, Tamar Harel, Shalini N. Jhangiani, Zeynep H. Coban Akdemir, Steven Buyske, Davut Pehlivan, Claudia M. B. Carvalho, Samantha Baxter, Nara Sobreira, Pengfei Liu, Nan Wu, Jill A. Rosenfeld, Sushant Kumar, Dimitri Avramopoulos, Janson J. White, Kimberly F. Doheny, P. Dane Witmer, Corinne Boehm, V. Reid Sutton, Donna M. Muzny, Eric Boerwinkle, Murat Günel, Deborah A. Nickerson, Shrikant Mane, Daniel G. MacArthur, Richard A. Gibbs, Ada Hamosh, Richard P. Lifton, Tara C. Matise, Heidi L. Rehm, Mark Gerstein, Michael J. Bamshad, David Valle, James R. Lupski, and Centers for Mendelian Genomics. Insights into genetics, human biology and disease gleaned from family based genomic studies. Genetics in Medicine, 21(4):798–812, Apr 2019. ISSN 1530-0366. doi: 10.1038/s41436-018-0408-7.

[2] Jennifer E. Posey. Genome sequencing and implications for rare disorders. Orphanet Journal of Rare Diseases, 14(1):153, Jun 2019. ISSN 1750-1172. doi: 10.1186/s13023-019-1127-0.

[3] Andries T. Marees, Hilde de Kluiver, Sven Stringer, Florence Vorspan, Emmanuel Curis, Cynthia Marie-Claire, and Eske M. Derks. A tutorial on conducting genome-wide association studies: Quality control and statistical analysis. International Journal of Methods in Psychiatric Research, 27(2):e1608, 2018. doi: https://doi.org/10.1002/mpr.1608.

[4] Stephen Turner, Loren L. Armstrong, Yuki Bradford, Christopher S. Carlson, Dana C. Crawford, Andrew T. Crenshaw, Mariza de Andrade, Kimberly F. Doheny, Jonathan L. Haines, Geoffrey Hayes, Gail Jarvik, Lan Jiang, Iftikhar J. Kullo, Rongling Li, Hua Ling, Teri A. Manolio, Martha Matsumoto, Catherine A. McCarty, Andrew N. McDavid, Daniel B. Mirel, Justin E. Paschall, Elizabeth W. Pugh, Luke V. Rasmussen, Russell A. Wilke, Rebecca L. Zuvich, and Marylyn D. Ritchie. Quality control procedures for genome-wide association studies. Current Protocols in Human Genetics, 68(1):1.19.1–1.19.18, 2011. doi: https://doi.org/10.1002/0471142905.hg0119s68.

[5] Monica D. Ramstetter, Thomas D. Dyer, Donna M. Lehman, Joanne E. Curran, Ravindranath Duggirala, John Blangero, Jason G. Mezey, and Amy L. Williams. Benchmarking relatedness inference methods with genome-wide data from thousands of relatives. Genetics, 207(1):75–82, 2017. ISSN 1943-2631. doi: 10.1534/genetics.117.1122.

[6] Cathie Sudlow, John Gallacher, Naomi Allen, Valerie Beral, Paul Burton, John Danesh, Paul Downey, Paul Elliott, Jane Green, Martin Landray, Bette Liu, Paul Matthews, Giok Ong, Jill Pell, Alan Silman, Alan Young, Tim Sprosen, Tim Peakman, and Rory Collins. UK Biobank: An Open Access Resource for Identifying the Causes of a Wide Range of Complex Diseases of Middle and Old Age. PLOS Medicine, 12(3):1–10, 03 2015. doi: 10.1371/journal.pmed.1001779.

[7] Alexander Gusev, Jennifer K. Lowe, Markus Stoffel, Mark J. Daly, David Altshuler, Jan L. Breslow, Jeffrey M. Friedman, and Itsik Pe’Er. Whole population, genome-wide mapping of hidden relatedness. Genome Research, 19(2):318–326, 2009. ISSN 1088-9051. doi: 10.1101/gr.081398.108.

[8] Daniel N. Seidman, Sushila A. Shenoy, Minsoo Kim, Ramya Babu, Ian G. Woods, Thomas D. Dyer, Donna M. Lehman, Joanne E. Curran, Ravindranath Duggirala, John Blangero, and Amy L. Williams. Rapid, Phase-free Detection of Long Identity-by-Descent Segments Enables Effective Relationship Classification. American Journal of Human Genetics, 106(4):453–466, 2020. ISSN 1537-6605. doi: 10.1016/j.ajhg.2020.02.012.

[9] Daniel N. Seidman, Sushila A. Shenoy, Minsoo Kim, Ramya Babu, Ian G. Woods, Thomas D. Dyer, Donna M. Lehman, Joanne E. Curran, Ravindranath Duggirala, John Blangero, and Amy L. Williams. Rapid, phase-free detection of long identity-by-descent segments enables effective relationship classification. The American Journal of Human Genetics, 106(4):453–466, 2020. ISSN 0002-9297. doi: https://doi.org/10.1016/j.ajhg.2020.02.012.

[10] William A. Freyman, Kimberly F. Mcmanus, Suyash S. Shringarpure, Ethan M. Jewett, Katarzyna Bryc, and Adam Auton. Fast and Robust Identity-by-Descent Inference with the Templated Positional Burrows-Wheeler Transform. Molecular Biology and Evolution, 38(5):2131–2151, 2021. ISSN 1537-1719. doi: 10.1093/molbev/msaa328.

[11] Monica D. Ramstetter, Sushila A. Shenoy, Thomas D. Dyer, Donna M. Lehman, Joanne E. Curran, Ravindranath Duggirala, John Blangero, Jason G. Mezey, and Amy L. Williams. Inferring identical-by-descent sharing of sample ancestors promotes high-resolution relative detection. The American Journal of Human Genetics, 103(1):30–44, Jul 2018. ISSN 0002-9297.

[12] Chad D. Huff, David J. Witherspoon, Tatum S. Simonson, Jinchuan Xing, W. Scott Watkins, Yuhua Zhang, Therese M. Tuohy, Deborah W. Neklason, Randall W. Burt, Stephen L. Guthery, Scott R. Woodward, and Lynn B. Jorde. Maximum-likelihood estimation of recent shared ancestry (ERSA). Genome Research, 21(5):768–774, 2011. ISSN 1088-9051. doi: 10.1101/gr.115972.110.

[13] Hong Li, Gustavo Glusman, Hao Hu Shankaracharya, Juan Caballero, Robert Hubley, David Witherspoon, Stephen L. Guthery, Denise E. Mauldin, Lynn B. Jorde, Leroy Hood, Jared C. Roach, and Chad D. Huff. Relationship Estimation from Whole-Genome Sequence Data. PLoS Genetics, 10(1), 2014. ISSN 1553-7390. doi: 10.1371/journal.pgen.1004144.

[14] Ani Manichaikul, Josyf C. Mychaleckyj, Stephen S. Rich, Kathy Daly, Michèle Sale, and Wei-Min Chen. Robust relationship inference in genome-wide association studies. Bioinformatics (Oxford, England), 26(22):2867–2873, Nov 2010. ISSN 1367-4811. doi: 10.1093/bioinformatics/btq559.

[15] Po-Ru Loh, Petr Danecek, Pier Francesco Palamara, Christian Fuchsberger, Yakir A Reshef, Hilary K Finucane, Sebastian Schoenherr, Lukas Forer, Shane McCarthy, Goncalo R. Abecasis, Richard Durbin, and Alkes L Price. Reference-based phasing using the Haplotype Reference Consortium panel. Nature Genetics, 48(11):1443–1448, Nov 2016. ISSN 1546-1718. doi: 10.1038/ng.3679.

[16] Christian Fuchsberger, Gonçalo R. Abecasis, and David A. Hinds. Minimac2: Faster genotype imputation. Bioinformatics, 31 (5):782–784, 2015. ISSN 1460-2059. doi: 10.1093/bioinformatics/btu704.

[17] F. Mölder, K. P. Jablonski, B. Letcher, M. B. Hall, C. H. Tomkins-Tinch, V. Sochat, J. Forster, S. Lee, S. O. Twardziok, A. Kanitz, A. Wilm, M. Holtgrewe, S. Rahmann, S. Nahnsen, and J. Köster. Sustainable data analysis with snakemake [version 2; peer review: 2 approved]. F1000Research, 10(33), 2021. doi: 10.12688/f1000research.29032.2.

[18] Anaconda software distribution, 2020. URL https://docs.anaconda.com/.

[19] Dirk Merkel. Docker: lightweight linux containers for consistent development and deployment. Linux journal, 2014(239):2, 2014.

[20] Task Execution Service (TES) API, 2022. URL https://github.com/ga4gh/task-execution-schemas.

[21] B. D. O’Connor, D. Yuen, V. Chung, A. G. Duncan, X. K. Liu, J. Patricia, B. Paten, L. Stein, and V. Ferretti. The Dockstore: enabling modular, community-focused sharing of Docker-based genomics tools and workflows [version 1; peer review: 2 approved]. F1000Research, 6(52), 2017. doi: 10.12688/f1000research.10137.1.

[22] Global Alliance for Genomics and Health (GA4GH), 2022. URL https://www.ga4gh.org/.

[23] Natalie A. Twine, Piotr Szul, Lyndal Henden, Emily P. McCann, Ian P. Blair, Kelly L. Williams, and Denis C. Bauer. TRIBES: A user-friendly pipeline for relatedness detection and disease gene discovery. bioRxiv, pages 0–1, 2019. doi: 10.1101/686253.

[24] Madison Caballero, Daniel N. Seidman, Ying Qiao, Jens Sannerud, Thomas D. Dyer, Donna M. Lehman, Joanne E. Curran, Ravindranath Duggirala, John Blangero, Shai Carmi, and Amy L. Williams. Crossover interference and sex-specific genetic maps shape identical by descent sharing in close relatives. PLoS Genetics, 15(12):1–29, 2019. ISSN 1553-7404. doi: 10.1371/journal.pgen.1007979.

[25] Claude Bherer, Christopher L. Campbell, and Adam Auton. Refined genetic maps reveal sexual dimorphism in human meiotic recombination at multiple scales. Nature Communications, 8, 2017. ISSN 2041-1723. doi: 10.1038/ncomms14994.

[26] Anders Albrechtsen, Ida Moltke, and Rasmus Nielsen. Natural selection and the distribution of identity-by-descent in the human genome. Genetics, 186(1):295–308, Sep 2010. ISSN 1943-2631. doi: 10.1534/genetics.110.113977.

[27] Catherine A. Ball, Mathew Barber, Jake K. Byrnes, Peter Carbonetto, Ken Chahine, Ross E. Curtis, Julie M. Granka, Eunjung Han, Eurie L. Hong, Amir R. Kermany, Natalie M. Myres, Keith Noto, Jianlong Qi, Kristin A. Rand, Yong Wang, and Lindsay Willmore. AncestryDNA Matching White Paper Discovering genetic matches across a massive, expanding genetic database. 2016.

[28] Peter J. Rousseeuw. Least median of squares regression. Journal of the American Statistical Association, 79(388):871–880, 1984. doi: 10.1080/01621459.1984.10477105.

[29] Funnel, 2022. URL https://github.com/ohsu-comp-bio/funnel.

[30] Allen Ancient DNA Resource (AADR), 2021. URL https://reichdata.hms.harvard.edu/pub/datasets/amh_repo/curated_releases/index_v44.3.html. Accessed: 2021-02-27.

[31] Doron M. Behar, Mait Metspalu, Yael Baran, Naama M. Kopelman, Bayazit Yunusbayev, Ariella Gladstein, Shay Tzur, Hovhannes Sahakyan, Ardeshir Bahmanimehr, Levon Yepiskoposyan, Kristiina Tambets, Elza K. Khusnutdinova, Alena Kushniarevich, Oleg Balanovsky, Elena Balanovsky, Lejla Kovacevic, Damir Marjanovic, Evelin Mohailov, Anastasia Kouvatsi, Costas Triantaphyllidis, Roy J. King, Ornella Semino, Antonio Torroni, Michael F. Hammer, Ene Metspalu, Karl Skorecki, Saharon Rosset, Eran Halperin, Richard Villems, and Noah A. Rosenberg. No Evidence from Genome-wide Data of a Khazar Origin for the Ashkenazi Jews. Human Biology, 85(6):859–900, 2013. ISSN 0018-7143, 1534-6617. doi: 10.13110/humanbiology.85.6.0859.

[32] Zhong Zhuang, Alexander Gusev, Judy Cho, and Itsik Pe’er. Detecting Identity by Descent and Homozygosity Mapping in Whole-Exome Sequencing Data. PLoS ONE, 7(10):1–7, 2012. ISSN 1932-6203. doi: 10.1371/journal.pone.0047618.

[33] Henn B. M., Hon L., Macpherson J. M., Eriksson N., Saxonov S., Pe’er I., and other. Cryptic distant relatives are common in both isolated and cosmopolitan genetic samples. PLoS ONE, 7(4), 2012. ISSN 1932-6203. doi: 10.1371/journal.pone.0034267.

[34] Atlas Biomed, 2022. URL https://atlasbiomed.com/uk.

[35] The 1000 Genomes Project Consortium. A global reference for human genetic variation. Nature, 526(7571):68–74, 2015. ISSN 1476-4687. doi: 10.1038/nature15393.

